# Oxidative stress induces release of mitochondrial DNA into the extracellular space in human placental villous trophoblast BeWo cells

**DOI:** 10.1101/2024.02.02.578433

**Authors:** Jennifer J. Gardner, Spencer C. Cushen, Reneé de Nazaré Oliveira da Silva, Jessica L. Bradshaw, Nataliia Hula, Isabelle K. Gorham, Selina M. Tucker, Zhengyang Zhou, Rebecca L. Cunningham, Nicole R. Phillips, Styliani Goulopoulou

## Abstract

Circulating cell-free mitochondrial DNA (ccf-mtDNA) is an indicator of cell death, inflammation, and oxidative stress. ccf-mtDNA differs in pregnancies with placental dysfunction from healthy pregnancies and the direction of this difference depends on gestational age and method of mtDNA quantification. Reactive oxygen species (ROS) trigger release of mtDNA from non-placental cells; yet it is unknown whether trophoblast cells release mtDNA in response to oxidative stress, a common feature of pregnancies with placental pathology. We hypothesized that oxidative stress would induce cell death and release of mtDNA from trophoblast cells. BeWo cells were treated with antimycin A (10-320 μM) or rotenone (0.2-50 μM) to induce oxidative stress. A multiplex real-time quantitative PCR (qPCR) assay was used to quantify mtDNA and nuclear DNA in membrane bound, non-membrane bound, and vesicular-bound forms in cell culture supernatants and cell lysates. Treatment with antimycin A increased ROS (p<0.0001), induced cell necrosis (p=0.0004) but not apoptosis (p=0.6471) and was positively associated with release of membrane-bound and non-membrane bound mtDNA (p<0.0001). Antimycin A increased mtDNA content in exosome-like extracellular vesicles (vesicular-bound form; p=0.0019) and reduced autophagy marker expression (LC3A/B, p=0.0002; p62, p<0.001). Rotenone treatment did not influence mtDNA release or cell death (p>0.05). Oxidative stress induces release of mtDNA into the extracellular space and causes non-apoptotic cell death and a reduction in autophagy markers in BeWo cells, an established *in vitro* model of human trophoblast cells. Intersection between autophagy and necrosis may mediate the release of mtDNA from the placenta in pregnancies exposed to oxidative stress.

**NEW & NOTEWORTHY:** This is the first study to test whether trophoblast cells release mitochondrial DNA in response to oxidative stress and to identify mechanisms of release and biological forms of mtDNA from this cellular type. This research identifies potential cellular mechanisms that can be used in future investigations to establish the source and biomarker potential of circulating mitochondrial DNA in preclinical experimental models and humans.

## INTRODUCTION

The placenta forms a critical interface between the mother and the developing fetus by functioning as the master regulator of fetal-maternal exchange of metabolic, inflammatory, and endocrine factors (1–3). The hemochorial orientation of the human placenta facilitates this communication by providing a limited cellular barrier between the maternal blood and fetal epithelium (4). This cellular barrier that forms the fetal-maternal interface is composed of trophoblast cells. In pathological conditions, trophoblast stress results in the release of proinflammatory and vasoactive factors that due to this hemochorial architecture can directly access the maternal vasculature, affecting its function and maternal outcomes during pregnancy (2).

Circulating cell-free mtDNA (ccf-mtDNA) is a damage-associated molecular pattern that is recognized by pattern recognition receptors of the innate immune system to elicit inflammation and facilitate immune system activation (5). Ccf-mtDNA increases with advancing gestational age in maternal sera of healthy pregnant individuals(6), potentially mirroring placental and fetal developmental trajectories and trophoblast cell turn over. In preeclampsia, a severe hypertensive disorder of pregnancy with placental pathology, ccf-mtDNA concentrations differ from healthy controls. Some studies have reported an increase (7, 8) and others showed a reduction(9, 10) in maternal ccf-mtDNA in patients with preeclampsia, depending on gestational age and method of mtDNA quantification (11).

In non-pregnant states, various types of cells have the ability to release mtDNA into circulation (12–14). In fact, damaged or dying non-placental cells release mtDNA into the extracellular space during programmed cell death (i.e., apoptosis) or passive cell death (i.e., necrosis) (5, 11, 15). An increase in production of reactive oxygen species (ROS) in non-placental tissues is a known trigger of mtDNA release (16, 17). In pregnancies with placental ischemia, a common feature of preeclampsia and a major contributor to adverse maternal and perinatal outcomes (2, 18), the placenta shows signs of necrosis, exaggerated apoptosis, and oxidative stress (2). Nevertheless, it remains unknown whether placental cells could also release mtDNA in response to activation of these mechanisms. Thus, the mechanisms of placental release of extracellular mtDNA and the ability of placental cells to contribute to ccf-mtDNA remain unknown.

The objective of this study was to elucidate the relationship between oxidative stress, cell death, and mtDNA release in trophoblast cells, which is the primary cellular unit of the maternal-fetal interface in the placenta. We hypothesized that trophoblast cells exposed to elevated levels of oxidative stress (i.e., reactive oxygen species, ROS) would release mtDNA into the extracellular space via cell-death dependent mechanisms. To test this hypothesis, we used a well-established *in vitro* model of human trophoblast cells (BeWo cells). Given that mtDNA is detected in the circulation in various biological forms (11), we measured membrane bound, non-membrane bound, and vesicular-bound mtDNA.

## MATERIALS AND METHODS

### Chemicals and Reagents

All chemicals and reagents are included in Supplemental Table 1 unless otherwise indicated.

### Cell culture

BeWo choriocarcinoma cells (sex: male, ATCC, cat# CCL-98) are a model of human epithelial cytotrophoblast cells (mononuclear) that form the progenitor trophoblast cells of the placenta. BeWo cells were grown in Kaighn’s Modification of Ham’s F-12 Medium (ATCC, cat# 30-2004), supplemented with 10% fetal bovine serum (FBS, ATCC, cat# 30-2020) and 1% penicillin/streptomycin (Gibco®, cat# 15140). The cells were maintained at 37°C in a > 90% relative humidity incubator with 5% CO_2_ (HeraCell 150i, Model No. 1335, Thermo Fisher Scientific). Routine subculturing was performed with 0.25% Trypsin-EDTA (Gibco®, cat# 25200-056). BeWo cells were authenticated through short tandem repeat profiling (ATCC). To establish the “trophoblast” nature of the BeWo cell line, cells were stained for anti-cytokeratin 7 using immunocytochemistry (Supplementary Methods & Supplementary Figure 1) (19).

### Experimental design and methods

After reaching 80% confluency, cells were treated for 4 hours with antimycin A (10, 50, 100, 320 μM; unless otherwise noted), or rotenone (0.2, 0.8, 5, 10, 25, 50 μM; unless otherwise noted) or their respective vehicles (ethanol (0.5% v/v) for antimycin A or DMSO (0.08% v/v for rotenone)) (20). Real-time qPCR with TaqMan™ probes and chemistry was used to quantify mtDNA and nuclear DNA in cell culture supernatants, cell lysates, and cell culture supernatant derived exosome-like extracellular vesicles (EV-BeWo; Figure 1). Oxidative stress was measured using a 2’,7’ –dichlorofluorescein diacetate (DCFDA) assay. Flow cytometry was used to assess cell viability, apoptosis, and necrosis. Western blot analysis was used to assess protein expression of autophagy markers.

**Figure 1.**
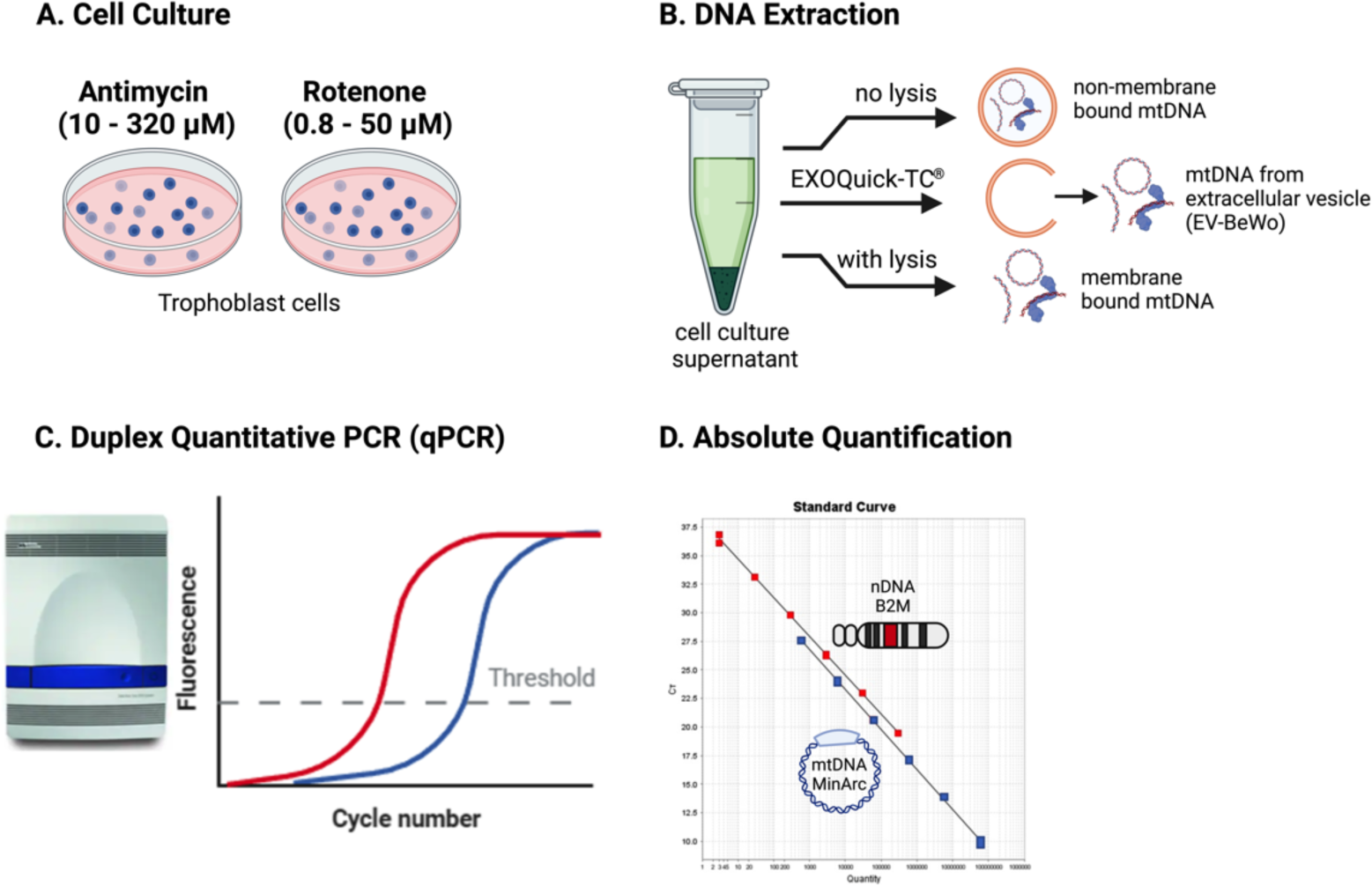
DNA extraction and quantification of mitochondrial DNA. (A) BeWo cells were exposed to rotenone or antimycin A for 4 hours. (B) DNA was extracted from the supernatant of BeWo cells with lysis buffer (membrane-bound form), without lysis buffer (non-membrane bound form), or with EXOQuick-TC^®^ method to isolate DNA stored within extravesicular membranes (EV-BeWo). (C) mtDNA and nDNA were quantified with a duplex qPCR protocol targeting (D) the nuclear primer (B2M) and the mtDNA primer (MinArc), using TaqMan© probes and chemistry. mtDNA, mitochondrial DNA; nDNA, nuclear DNA. Figure was made with Biorender.

### Isolation and characterization of exosome-like extracellular vesicles

FBS was depleted of exosomes via ultracentrifugation (Thermo Scientific, Sorvall WX + Ultra Series) at 10,000 xg for 2 hours as previously described(21). The exosome-depleted FBS aqueous layer was then filtered through a 0.2 μm-sized 50-mL tube top vacuum filter system (cat# 430320, Corning, NY). Exosome-depleted media (Kaighn’s Modification of Ham’s F-12 Medium supplemented with 10% exosome-depleted FBS and 1% penicillin/streptomycin) was used to seed BeWo cells into 100 mm petri dish at a density of 100,000 cell/ml. The cells were incubated for approximately 24 hours at 37°C until approximately 70% confluent. The cells were washed with phosphate-buffered saline (PBS) and then treated with antimycin A (320 μM) or vehicle for 4 hours. Extracellular vesicles were isolated from 10 mL of cell culture supernatant using the ExoQuick-TC^®^ Exosome Precipitation Solution (cat# EXOTC10A-1; System Biosciences) according to the manufacturer’s instructions with a few modifications. Supernatants were collected and centrifuged at 3000 xg for 15 minutes and then again at 7500 rpm for 10 minutes at 4°C to remove any cells and/or cellular debris. We added 1.5 times the recommended volume of ExoQuick-TC^®^ to 9.4 mL of clarified BeWo cell culture supernatant and incubated overnight. Extracellular vesicles were collected by centrifuging at 1500 xg for 30 minutes and removing the supernatant. This centrifugation step was repeated twice to ensure the complete removal of media from the pellet. Extracellular vesicles were then resuspended in 250 μl PBS and stored at −80°C until DNA extraction. Exosome-like extracellular vesicles from BeWo cells were designated as “EV-BeWo”.

Characterization of extracellular vesicles was performed by Systems Biosciences using an Exo-Check^TM^ exosome antibody array (ExoRay200B, cat# CSEQ100A-1, Palo Alto, CA) and nanoparticle tracking analysis (NTA, Nanosight, Version 2.3, Build 2.3.5.0033.7-Beta-7, cat# CSNANO100A-1). The array was exposed to 50 µg of exosome proteins isolated from supernatants of BeWo cells cultured in exosome-depleted media. The following exosome markers were used in the array: CD63 and CD81 (tetraspanins), EpCam (epithelial cell adhesion molecule; often found in cancer-derived exosomes), ANX5 (annexin 5), TSG01 (tumor susceptibility gene 101), FLOT-1 (flotillin-1), ICAM-1 (intercellular adhesion molecule 1), ALIX (programmed cell death 6 interacting protein (PDCD6IP). The NTA analysis with Systems Biosciences used a proprietary fluorescence dye that binds to intact membrane of extracellular vesicles, ExoGlow-NTA, to detect size and concentration of extracellular vesicles.

### DNA extraction

DNA was measured in cell suspensions and cell culture supernatants from the same experiment. Cell culture supernatants were collected from cell culture flask following 4-hour exposure to rotenone, antimycin A, or vehicle. Adherent cells were collected using 0.25% trypsin-EDTA and centrifuging for 5 minutes at 400 xg at 4°C. Cell pellets were resuspended in elution buffer. Independent experiments were performed to determine DNA content in extracellular vesicles.

#### Cell Suspensions

DNA was extracted from cell suspensions using the Mag-Bind® Blood & Tissue DNA HDQ extraction kit (Omega Bio-Tek, Nocross, Georgia, USA) according to manufacturer’s specifications in the Mag-Bind® Blood protocol. DNA extracts were quantified using the Qubit dsDNA Broad Range assay kit (ThermoFisher Scientific) according to manufacturer’s recommendations. Samples with low concentrations were concentrated by spinning in the Vacufuge vacuum concentrator (Eppendorf) until concentrations reached at least 2.5 ng/µL adequate for use in qPCR.

#### Cell Culture Supernatants

DNA was extracted from supernatants twice using the Mag-Bind® Blood & Tissue DNA HDQ extraction kit (Omega Bio-Tek) according to manufacturer’s specifications in the Mag-Bind® Blood protocol with modifications. The first of the two DNA extractions were performed including a lysis step and the second extraction did not include the lysis step as previously published (10) to determine the contribution of membrane bound and non-membrane bound DNA, respectively.

#### Extracellular Vesicles

DNA was extracted from 200 µL of the resuspended extracellular vesicles using the DNeasy Blood & Tissue Kit (Qiagen, cat# 69504) according to the instructions in the Purification of Total DNA from Animal Blood or Cells (Spin-Column Protocol). DNA samples were eluted using 200 µL nuclease-free water instead of the elution buffer (Buffer AE) provided in the extraction kit. After DNA extraction, samples were concentrated by spinning in the Vacufuge vacuum concentrator (Eppendorf) until they reached a volume of 20 µL.

### DNA quantification via polymerase chain reaction (qPCR)

Multiplex quantification of nuclear DNA and mtDNA was performed according to Phillips et al., 2014 (22) and as we previously published (10), with some modifications. Namely, the deletion target was omitted. The master mix was modified as follows: 2 µL of each primer MinArc (0.625 µM) and B2M (7.5 µM); 1.25 µL of each probe MinArc (2 µM) and B2M (4 µM); 12.5 µL of TaqMan Universal MasterMix II, no UNG (Applied Biosystems, cat# 4440040); 2 µL of DNA extract. The concentration of DNA obtained from the supernatants was too low to be quantified, so it was used directly for qPCR without quantifying. DNA (5 ng) from cell suspension extracts were added to qPCR reaction mix. Standard curves based on known synthetic DNA copy number for both targets were used. R^2^ values and amplification efficiencies of the standard curves were analyzed for each plate to verify repeatability and monitor for potential batch effects. Negative controls were included on every run to monitor basal off-target mtDNA amplification.

### Cytosolic oxidative stress measurements via DCFDA assay

BeWo cells were seeded into a black-walled clear bottom 96-well plate at a density of 25,000 cells/well and grown until 80% confluent. Cells were washed with 100 μl PBS and stained with 25 μM of DCFDA for 45 minutes in a cell culture incubator (37°C, 5% CO_2_) before treatment with antimycin A, rotenone, or their respective vehicles in phenol free F-12K media. Following a 4-hour incubation period with antimycin A or rotenone, cells were analyzed on a fluorescent plate reader (Biotek Synergy HTX, Agilent Technologies, Inc., Santa Clara, CA) at excitation of 485/20 nm and emission 528/20 nm. Cell treatments were assayed in duplicate, and fluorescence intensity averaged from five independent experiments. Data presented as fluorescence intensity after background (blank wells with media only) subtraction.

### Flow cytometry

A cell-based flow cytometry assay kit (Abcam, cat# ab176749) was used to simultaneously detect apoptotic, necrotic, and viable BeWo cells after treatment with antimycin A or rotenone. For these experiments, BeWo cells were plated onto 60 mm polystyrene dishes and grown until 80% confluent. Cells were treated with antimycin A (10, 50, 100, 320 µM) or rotenone (0.8, 5, 10, 25, 50 µM) as described above, and adherent cells were collected using 0.25% trypsin-EDTA and centrifuged for 5 minutes at 400 xg at 4°C. Pellets were rinsed with ice-cold PBS and centrifuged for 5 minutes at 200 xg at 4°C. Cells were resuspended in 200 μL of assay buffer, according to the manufacturer’s instructions and then stained for 30 minutes at room temperature with 2 μL of 7-AAD (Ex/Em: 546/647 nm), apopoxin (Ex/Em: 490/525 nm), and cyto-calcein violet (Ex/Em: 405/450 nm) to detect necrosis, apoptosis, and viability, respectively. Dye concentration was determined after preliminary titration experiments. After staining, samples were then read using an LSR II FACS/flow cytometer (BD Biosciences; San Jose, CA) and analyzed using FlowJo flow cytometry software (v.10; Becton, Dickinson & Company; Ashland, OR). Viable cells were defined as cells stained positive for cyto-calcein but negative for 7AAD and Apopoxin (CC^+^, 7ADD^-^, Apo^-^). Apoptotic cells were defined as cells stained positive for Apopoxin (Apo^+^, 7AAD^+^ and Apo^+^, 7AAD^-^), while necrotic cells were defined as those cells stained positive for 7AAD and negative for Apopoxin (7AAD^+^, Apo^-^). We identified two populations of cells. Each gating strategy was applied separately to each population (Supplementary Figure 2A, 2C). We calculated the sum of stained cells from population 1 and population 2 then divided by the total cell count from both populations to determine the percentage of viability, apoptosis, and necrosis (Supplementary Figure 2B, 2D, 2E).

### Western Blots

To further understand mechanisms associated with the release of mtDNA, we focused on experiments with antimycin A. BeWo cells were seeded into 6-well plates at a density of 100,000 cells/ml and grown until 80% confluent. Trophoblast cells were treated with antimycin A (10, 100, or 320 μM) or vehicle for four hours. Protein was extracted with lysis buffer containing cOmplete Lysis-M with protease inhibitor cocktail tablets and protein concentrations were determined using Pierce BCA Protein Assay Samples were denatured with ß-mercaptoethanol and heated to 100 °C for 5 minutes. Equal amounts of protein (20 μg) were loaded on to 10% or 15% SDS-PAGE gels for 3 hours at 100 volts. Proteins were transferred for 90 minutes to nitrocellulose or polyvinylidene difluoride (PVDF) membranes. Afterwards, membranes were blocked for 1 hour with 3% Bovine Serum Albumin or with 5% non-fat dry milk (for LC3 A/B) as described by Lima et al. (23) Primary antibodies were dissolved with 3% BSA in tris buffered saline (TBST) and membranes were incubated overnight at 4°C with SQSTM1/p62 (Cell Signaling, cat# 88588, 1:1000). The primary LC3 A/B (Cell Signaling, cat# 4108s, 1:1000) was dissolved with 3% non-fat dry milk. Membranes were washed using TBST then incubated with secondary antibodies ((anti-rabbit IgG (H+L) DyLight 680 conjugate (LI-COR Biosciences, cat# 925-68071, 1:5000), anti-mouse IgG (H+L) DyLight 800 conjugate (LI-COR Biosciences, cat# 926-32210, 1:10000) and rabbit conjugated to horseradish peroxidase (HRP) (Cell Signaling, cat# 7074, 1:4000)), for 1 hour at room temperature. Data were acquired with Odyssey CLx LI-COR (LI-COR Biosciences) and analyzed with Image Studio v. 5.2 (LI-COR Biosciences). For LC3 A/B, data were obtained with SuperSignal^TM^ West Dura Extended Duration Substrate system (Thermo Fisher Scientific) and Azure 300 chemiluminescence detection system (Biosystem, USA), and band intensities were determined with ImageJ software 13.0.6 (NIH, USA). Values were normalized to total protein using ponceau staining. Representative immunoblots are illustrated in Supplementary Figure 3.

### Statistical analysis

Linear regression models were estimated to examine the association between each drug concentration (rotenone or antimycin A) and DNA quantities (membrane-bound and non-membrane bound mtDNA and nuclear DNA). Analyses were stratified by nuclear DNA (B2M) or mtDNA (MinArc) and adjusted for batch effects by including batch as a covariate. A DNA quantity value was considered as an outlier if it was 1.5 times interquartile range above the third quartile or below the first quartile and was then removed from analyses. Partial (Cohen’s) f^2^ were calculated to measure effect sizes. A partial f^2^ of 0.02, 0.15, and 0.35 represents a small, medium, and large effect, respectively (24).

One-way ANOVA with Dunnett’s multiple comparison test (for parametric data) or Kruskal-Wallis with Dunn’s multiple comparison test (for non-parametric data) were used to examine the effect of treatment on ROS activity (DCFD data) and cell death (flow cytometry data). Unpaired *t*-tests were used for group comparisons in EV-BeWo mtDNA content. Normality test (Shapiro-Wilk test), outlier removal (ROUT), and univariate analysis were performed using Prism (Version 8, GraphPad, San Diego, CA). Data are presented as mean ± SD (unless otherwise specified) and individual data points are included in figures. The significance level was set at ⍺= 0.05 for all comparisons and exact p values are presented.

## RESULTS

### Oxidative stress and mtDNA release from trophoblast cells

Cytosolic ROS activity was increased in response to antimycin A (one-way ANOVA, p<0.0001, Figure 2A) but not in response to rotenone (one-way ANOVA, p=1.79, Figure 2B).

**Figure 2.**
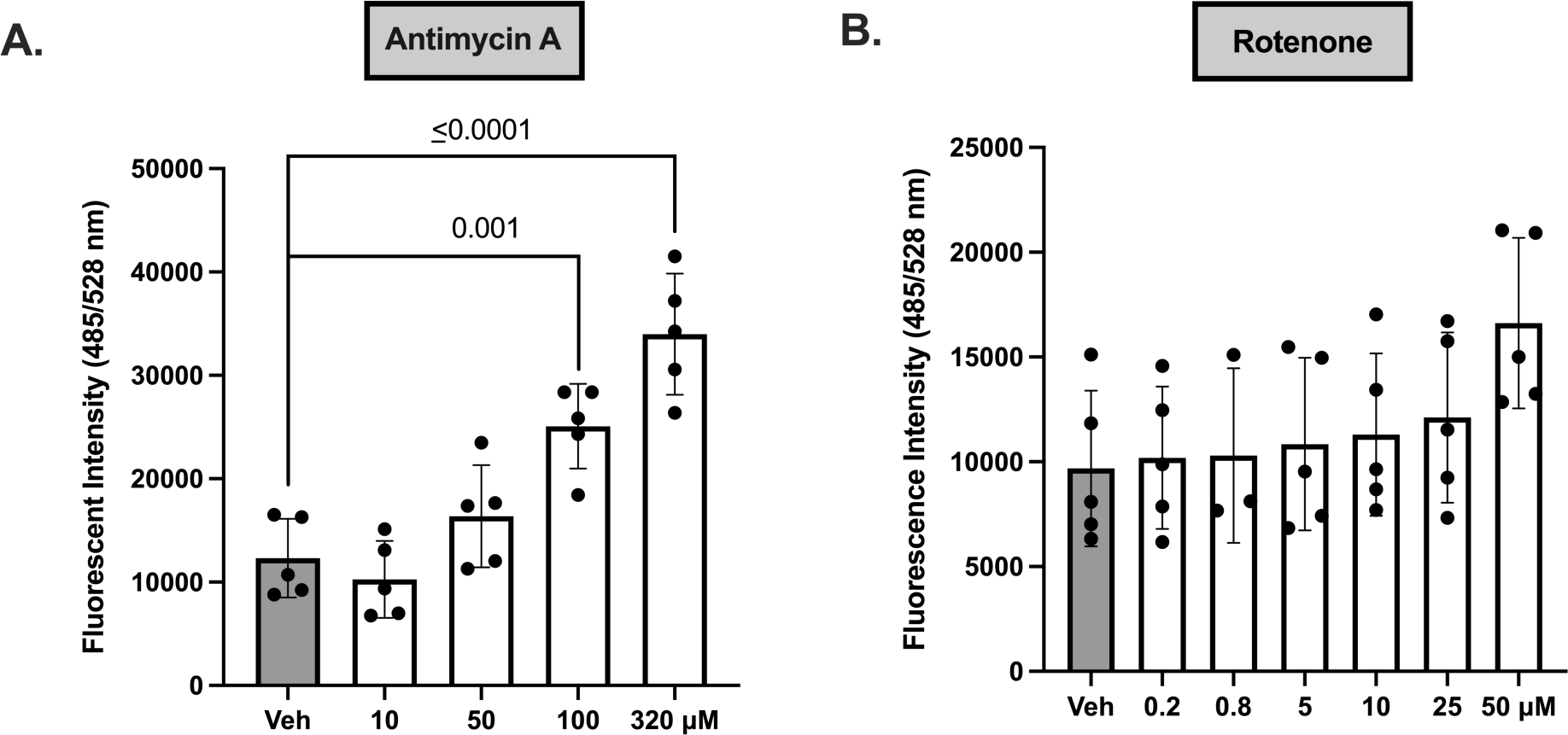
Cytosolic ROS levels in BeWo trophoblast cells following exposure to antimycin A or rotenone. A DCFDA assay was performed to determine ROS levels in BeWo cells treated with (A) antimycin A (n=5/concentration) or (B) rotenone (n=3-5/concentration). Significance was determined using a one-way ANOVA with Dunnett’s post hoc analysis. All values presented as means ± SD. n indicates independent experiments. DCFDA, 2’-7’-Dichlorodihydrofluorescein diacetate; ROS, reactive oxygen species.

There was a positive association between treatment with antimycin A and release of membrane-bound mtDNA (linear regression, p< 0.0001, partial f^2^=1.73, Figure 3A) and nuclear DNA (linear regression, p=0.024, partial f^2^=0.14, Figure 3B) in cell culture supernatants. Antimycin A treatment was not associated with intracellular mtDNA copy number (linear regression, p=0.62, partial f^2^=0.006, Figure 3C). There was no association between rotenone treatment and membrane-bound mtDNA (linear regression, p=0.43, partial f^2^=0.014, Figure 3D), nuclear DNA (linear regression, p=0.82, partial f^2^=0.001, Figure 3E) or intracellular mtDNA copy number (linear regression, p=0.054, partial f^2^=0.0849, Figure 3F).

**Figure 3.**
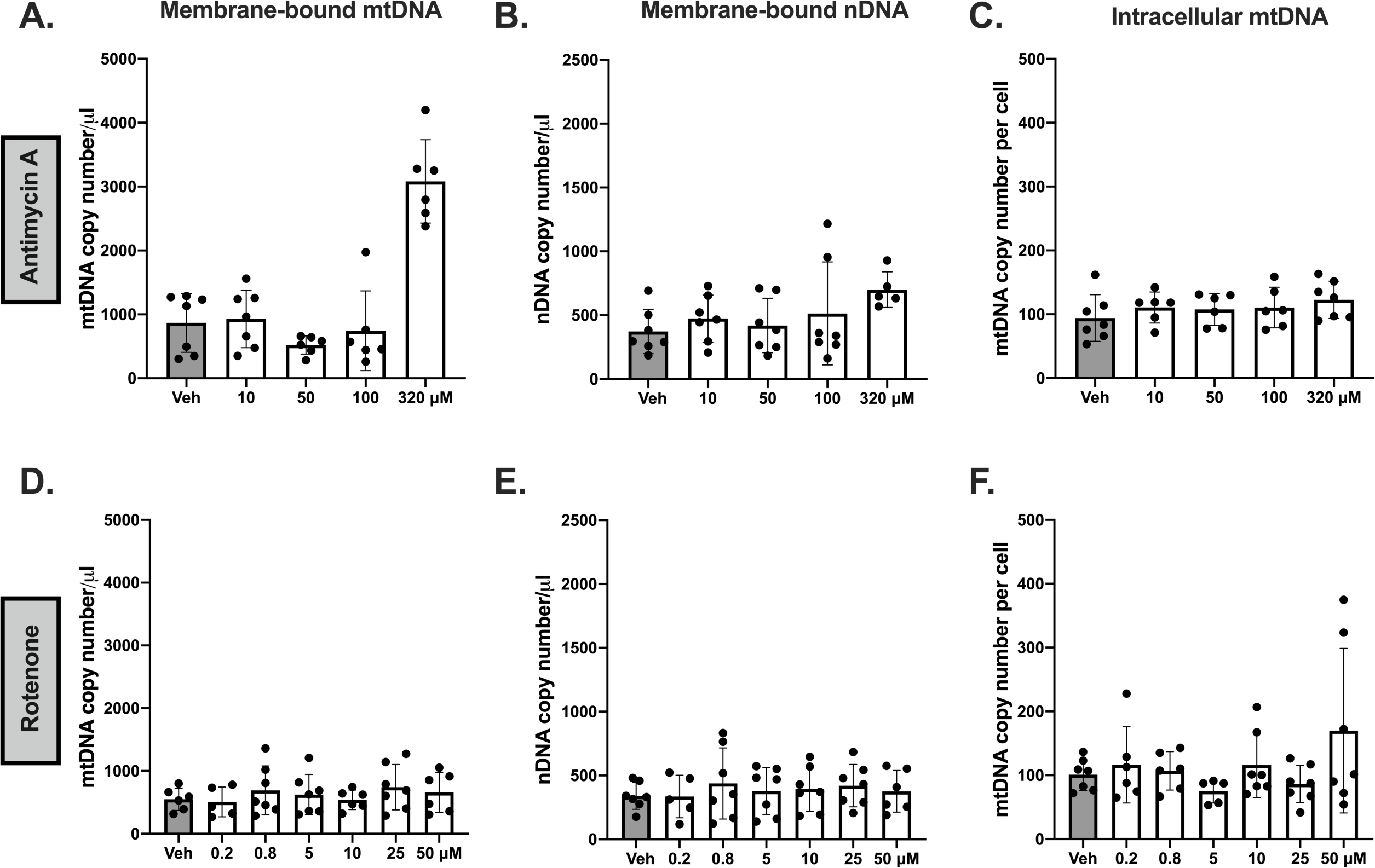
Membrane-bound extracellular mitochondrial (mtDNA) and nuclear DNA (nDNA) and intracellular mtDNA content in BeWo trophoblast cells in response to antimycin A and rotenone. Antimycin A increased the release of (A) membrane-bound mtDNA (p<0.0001) and (B) nuclear DNA (p=0.024), but there was no effect on (C) intracellular mtDNA copy number (p=0.62). Rotenone did not affect the release of (D) membrane-bound mtDNA (p=0.43), (E) nuclear DNA (p=0.82), or (F) intracellular mtDNA (p=0.054). All values presented as means ± SD. n=5-7 independent observations. Significance was determined using a linear regression model (drug concentration vs DNA quantities) stratified by nuclear DNA (B2M) or mtDNA (MinArc) and adjusted for batch effects.

Treatment with antimycin A was positively associated with non-membrane bound mtDNA in cell culture supernatants (linear regression, p<0.001, partial f^2^=0.84, Figure 4A), but it was not associated with non-membrane bound nuclear DNA (linear regression, p=0.051, partial f^2^=0.113, Figure 4B). Exposure to rotenone was not associated with non-membrane bound mtDNA (linear regression, p=0.180, partial f^2^=0.039, Figure 4C) or nuclear DNA (linear regression, p=0.115, partial f^2^=0.057, Figure 4D) in BeWo cell culture supernatants.

**Figure 4.**
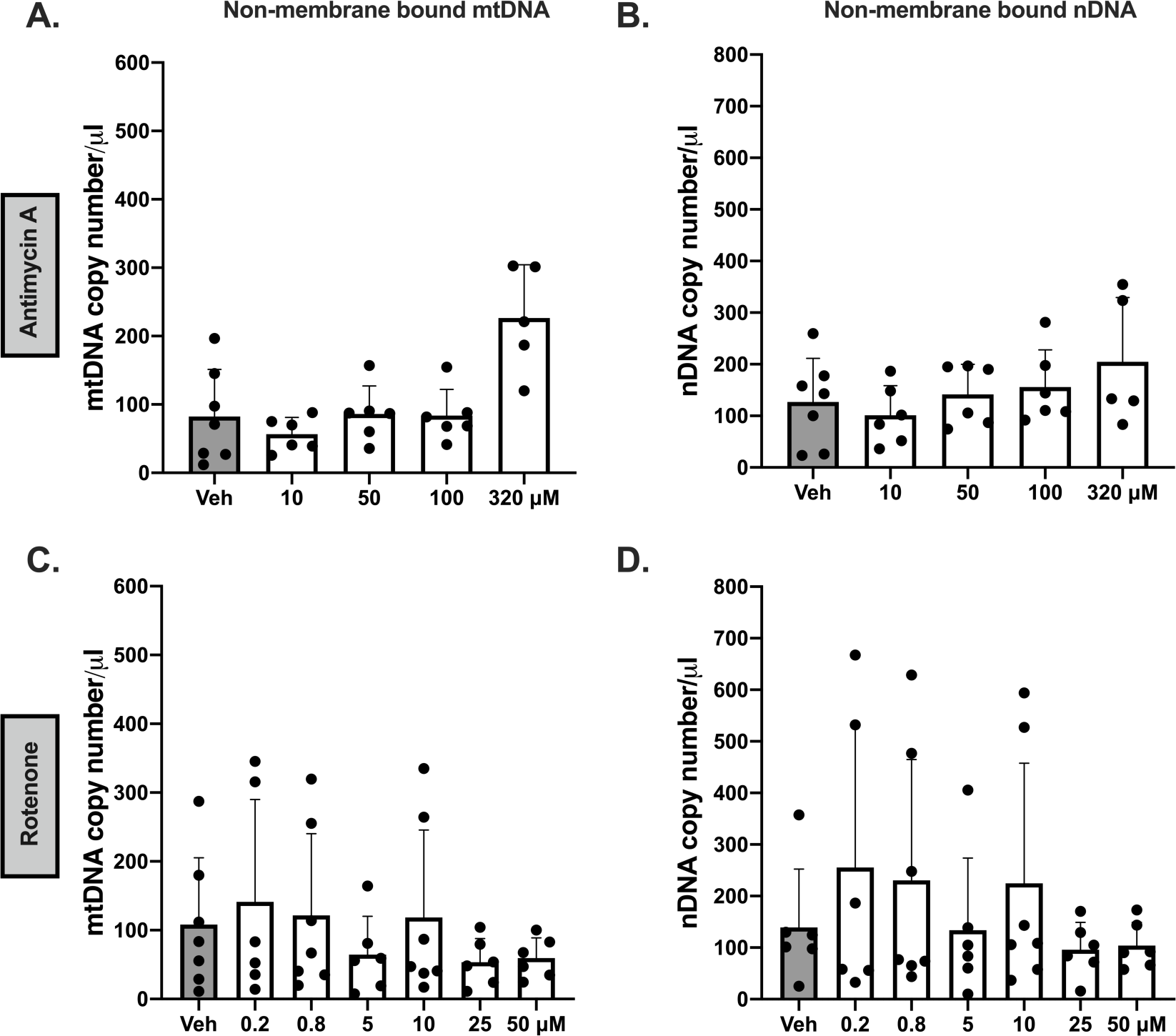
Non-membrane bound extracellular mitochondrial (mtDNA) and nuclear DNA (nDNA) in BeWo trophoblast cells in response to antimycin A and rotenone. Antimycin A increased the release of (A) non-membrane bound mtDNA (p< 0.001), but not (B) nDNA (p=0.051). Rotenone did not affect the release of (C) non-membrane bound mtDNA (p=0.180) or D) nuclear DNA (p=0.115). All values presented as means ± SD. n=5-7 independent observations. Significance was determined using a linear regression model (drug concentration vs DNA quantities) stratified by nuclear DNA (B2M) or mtDNA (MinArc) and adjusted for batch effects.

Antimycin A increased the membrane-bound mtDNA to nuclear DNA ratio (one-way ANOVA, p=0.016; Supplementary Figure 4A) and the non-membrane bound mtDNA to nuclear DNA ratio (one-way ANOVA, p=0.004; Supplementary Figure 4B). Rotenone did not affect the membrane-bound (Kruskal-Wallis, p=0.643) and non-membrane bound (one-way ANOVA, p>0.99) mtDNA to nuclear DNA ratio (Supplementary Figure 4C-D).

To determine if increases in membrane-bound DNA reflect an increase in vesicle-contained DNA content, we isolated EV-BeWo from supernatants of cells treated with antimycin A (320 µM) or vehicle. We chose this concentration of antimycin A because we previously observed an increase in membrane-bound mtDNA in response to this concentration but not in other concentrations. Antimycin A increased EV-BeWo contained mtDNA content (Unpaired *t*-test, p=0.0019, Figure 5A), nDNA (Unpaired *t*-test, p=0.0267, Figure 5B), and the mtDNA to nuclear DNA ratio (Unpaired *t*-test, p= 0.0581, Figure 5C) compared to vehicle. Semi-quantitative ExoCheck Exosome Antibody Array showed the isolated vesicles expressed exosome-associated protein markers (Figure 5D). Finite track length analysis of EV-BeWo revealed a size distribution of 117.1±0.9 nm (antimycin A, Figure 5E) and 75.2±1.0 nm (vehicle, Figure 5F).

**Figure 5.**
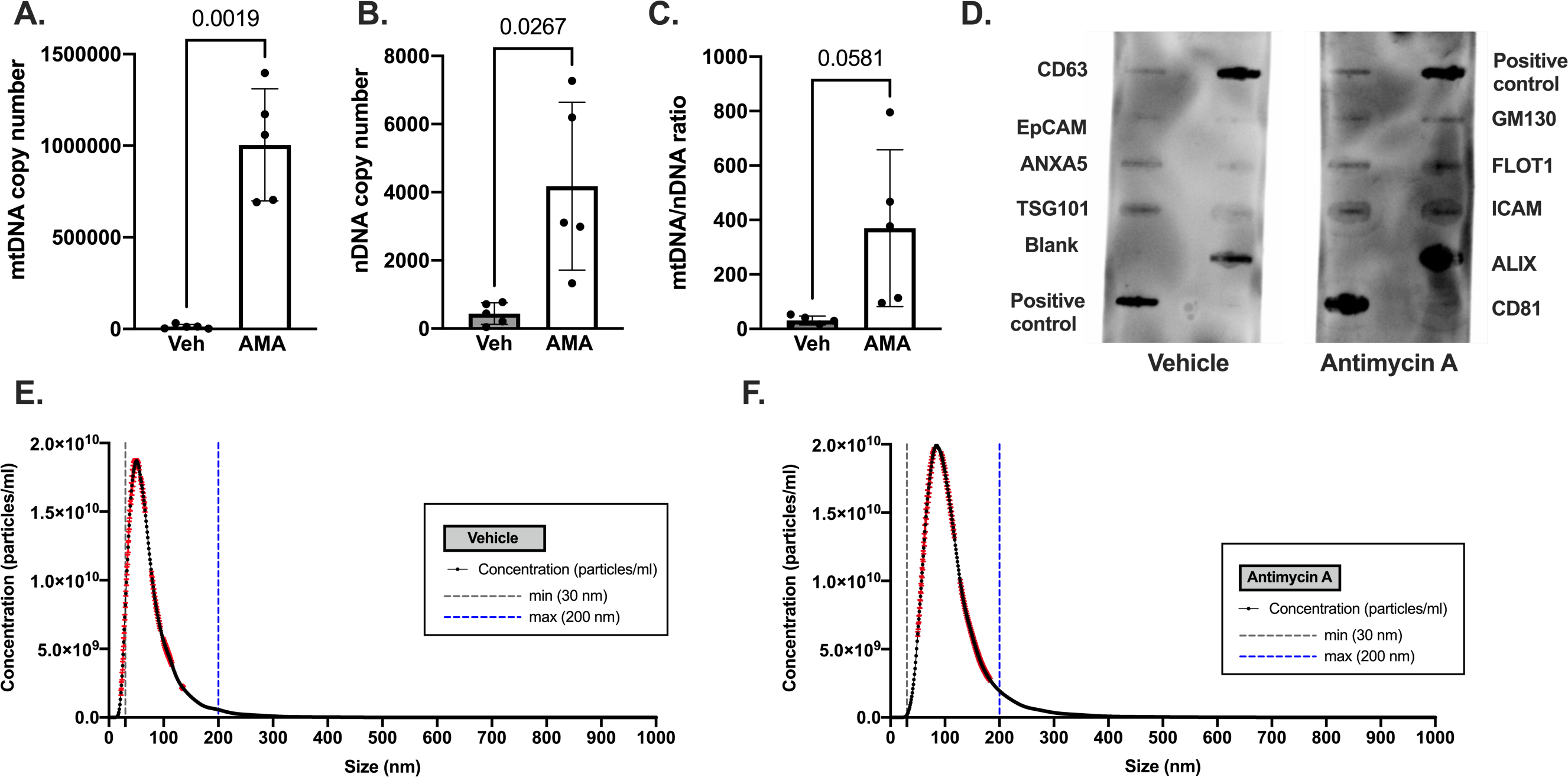
Mitochondrial DNA (mtDNA) and nuclear DNA (nDNA) content in BeWo-derived extracellular vesicles (EV-BeWo) in response to oxidative stress. EV-BeWo released in response to antimycin A had greater content of (A) mtDNA, (B) nuclear DNA, (C) mtDNA/nDNA ratio. All values presented as means ± SD. n=5 independent observations/treatment. Significance was determined using unpaired t-tests. (D) Semi-quantitative ExoCheck Exosome Antibody Array was exposed to 50 μg of exosomal proteins isolated from pooled BeWo cell supernatant using ExoQuick®. Positive exosomal markers include: CD63, CD81, ALIX, FLOT1, ICAM1, EpCam, ANXA5, and TSG101. A labeled positive control for HRP detection and a blank spot as a background control have been included. (E-F) Nanoparticle tracking analysis was used to determine size distribution (nm) of EVs from BeWo cells treated with (E) Vehicle and (F) antimycin A. Each graph depicts the average of 3 readings for each group. The threshold lines at 30 nm (grey dotted line) and 200 nm (blue dotted line) denote the size range of small EVs. Red error bars indicate ± SEM.

### Oxidative stress, cell death, and autophagy in trophoblast cells

Antimycin A decreased cell viability (one-way ANOVA, p<0.0001, Figure 6A), increased necrosis (one-way ANOVA, p=0.0004, Figure 6B), but it did not affect apoptosis (Kruskal Wallis, p= 0.65, Figure 6C). Rotenone had no effect on cell viability, necrosis, or apoptosis (p≥0.75, Figure 6D-F). Supplementary Figure 2 shows the gating strategy for the flow cytometry data analysis.

**Figure 6.**
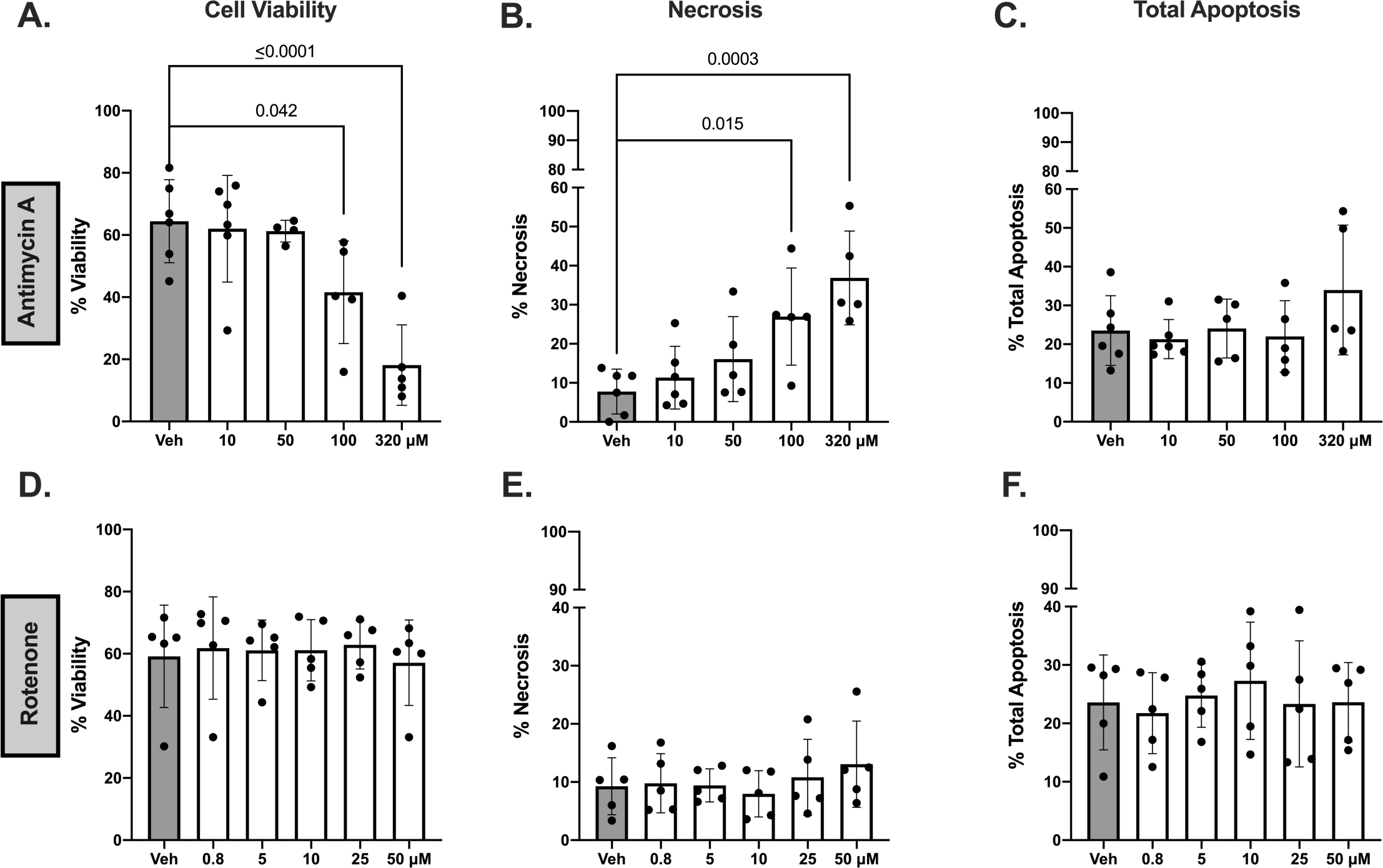
Effect of antimycin A and rotenone on cell viability and death in BeWo trophoblast cells. Antimycin A (n=5) (A) decreased cell viability, (B) increased necrosis, and (C) did not affect apoptosis. Rotenone (n=3-5) did not affect (D) cell viability, (E) necrosis, or (F) apoptosis. Data were analyzed using a one-way ANOVA with Dunnett’s multiple comparison test (for parametric data) or Kruskal-Wallis with Dunn’s multiple comparison test (for non-parametric). All cells are presented as a percentage of total cells. All values presented as means ± SD. n indicates independent observations/drug concentration.

Antimycin A reduced protein content of p62 (one-way ANOVA, p<0.0001, Figure 7A) and increased protein content of LC3 A/B II/I ratio (one-way ANOVA, p=0.0040, Figure 7B).

**Figure 7.**
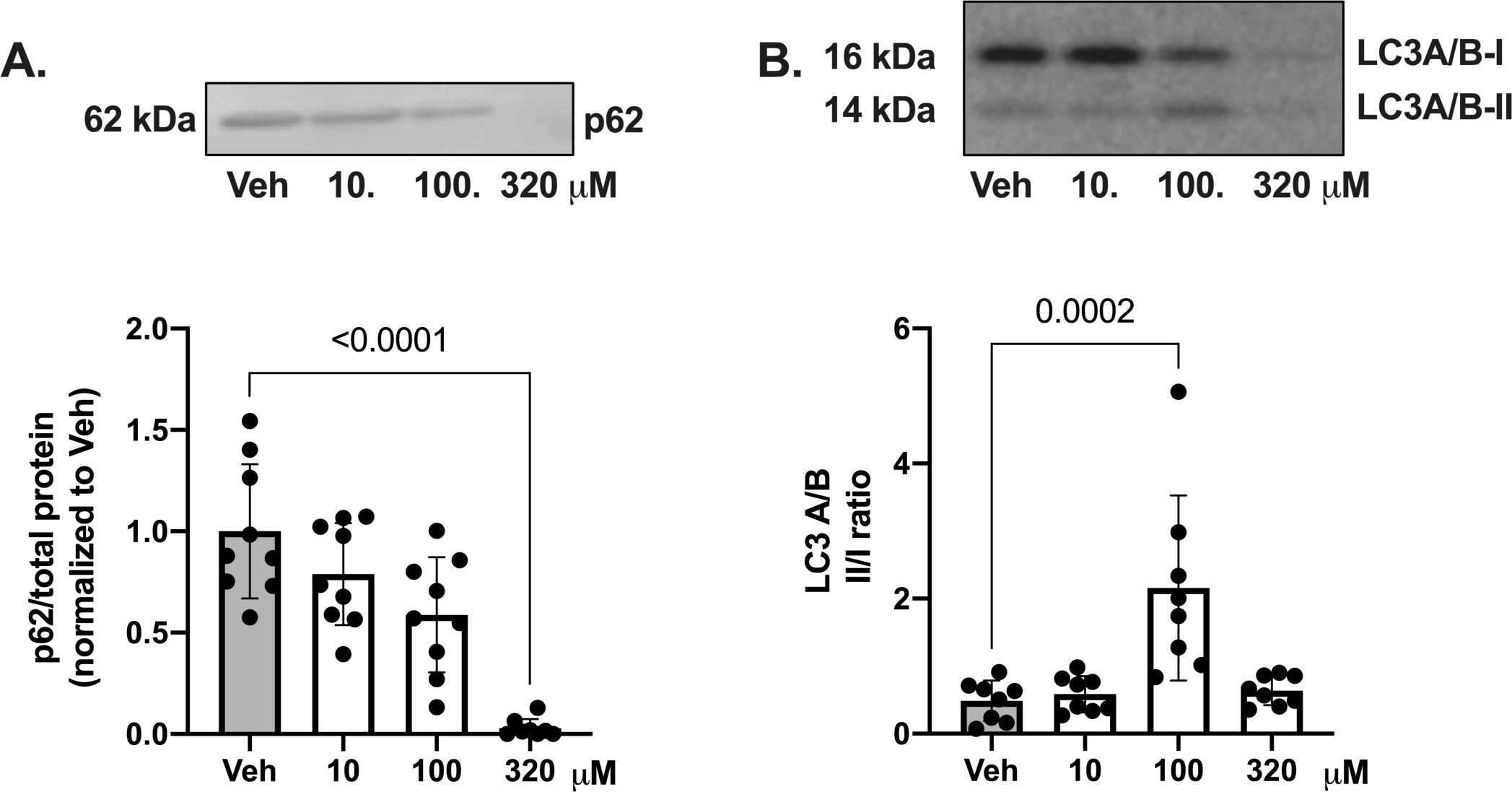
BeWo trophoblast cell protein content of autophagy markers p62 and LC3 following exposure to antimycin A. Antimycin A decreased (A) protein content of p62 and increased (B) LC3 A/B II/I ratio in BeWo cells. Representative immunoblots are presented above the graphs. Band intensity for p62 is normalized to total protein (Ponceau S stain). P62 summary data are presented relative to the mean of vehicle controls (mean value was set to 1). Values are shown as means ± SD. n= 7-9/drug concentration. Data were analyzed with one way ANOVA (and Dunnett’s multiple comparison test) or Kruskal-Wallis (and Dunn’s multiple comparison test).

## DISCUSSION

The main findings of this study are a) an increase in ROS triggers release of mtDNA in BeWo cells, a well-established *in vitro* model of human trophoblast cells and b) mtDNA is released from BeWo trophoblast cells in various biological forms, namely membrane bound, non-membrane bound, and vesicular bound. Furthermore, our results suggest that autophagy activation and cell necrosis, but not cell apoptosis, are involved in ROS-mediated mtDNA release from BeWo cells. This is the first study to interrogate the ability of trophoblast cells to release mtDNA in response to ROS, and to identify mechanisms of release and biological forms of mtDNA that derives from trophoblast cells.

ccf-mtDNA is used as an accessible blood-based surrogate of cellular stress and inflammation in order to assess disease progression and severity(12), survival rates(13), and disease prognosis (14) in non-obstetric pathologies. Establishing the cellular origin of ccf-mtDNA in the circulation may provide information about organ function, mechanisms of release, and cellular signaling (11). Various types of non-placental cells have the ability to release mtDNA including platelets (25), leukocytes (26), endothelial cells (27), and immortalized cell lines (27–32). We and others have shown that maternal ccf-mtDNA concentrations are dysregulated in obstetric complications such as preeclampsia and intrauterine growth restriction (7–10). The source of maternal ccf-mtDNA in these studies was not investigated and it was hypothesized that it may originate from the placenta and functions as an indicator of placental cell turnover and tissue function (7–10, 33). Indeed, recent studies confirm that first trimester human placental explants can release mtDNA (34). Similarly, our present findings demonstrate that BeWo cells, an *in vitro* model of trophoblast cells, release mtDNA constitutively and this release is exaggerated when the cells are exposed to stress.

An increase in ROS is a known trigger of mtDNA release in non-placental tissues (16, 17). Our data are the first to show that an increase in ROS triggers the release of mtDNA from BeWo trophoblast cells as well. Placental ROS is highly relevant to obstetric complications because it is associated with placental ischemia, mitochondrial dysfunction, and inflammation, all of which contribute to adverse perinatal outcomes (2, 35, 36). Exposure to pharmacological inhibition of the electron transport chain, either with rotenone or antimycin A (inhibitors of complexes I and III, respectively) or other cytotoxic agents such as hydrogen peroxide and low levels of oxygen are often used to induce ROS production in *in vitro* experimental set ups (37). Findings from these studies suggest an association between increased ROS production and cell death in placental (20, 38, 39) and non-placental cells (40, 41). Here, we posit that this relationship also mediates the release of mtDNA into the extracellular space in trophoblast cell cultures. Our data shows that an increase in trophoblast cell ROS triggers necrosis, but not apoptotic cell death, and causes a dose-dependent increase in the release of mtDNA. In the placental cell lines, BeWo, JEG-3, and Swan-71, exposure to rotenone or antimycin A led to a dose-dependent increase in ROS, and also led to apoptotic cell death in rotenone treated cells after 4 hours(20). ROS-mediated increase in apoptotic cell death has also been reported in response to hypoxia or hydrogen peroxide (38, 39). Non-apoptotic cell death and release of mtDNA, however, were not determined in any of these previous investigations. BeWo cells spontaneously form syncytia and the level of syncytialization could affect the sensitivity of the cells to ROS-mediated apoptosis (42). Hence, differences in levels of syncytialization before treatment between studies could explain differential outcomes in cell death (42, 43). Furthermore, disparate effects of rotenone and antimycin A on cell differentiation could explain an increased tolerance to rotenone in BeWo cells as indicated by reduced levels of ROS production and higher viability as compared to antimycin A (44, 45). Overall, our findings suggest that treatment type and ROS severity may contribute to the differences observed in cell death outcomes and mtDNA release in response to rotenone or antimycin A in BeWo cells (46).

ccf-mtDNA can be released in multiple biological forms under physiologic and pathologic conditions (47). In line with these findings, we demonstrate that mtDNA is released from trophoblast cells constitutively and in response to stress in both membrane-bound and non-membrane bound forms. These findings agree with our previous reports in human pregnancies complicated with preeclampsia (10). It has been suggested that membrane-bound form of mtDNA may be protected from degradation and play a role in signaling mechanisms primarily through extracellular vesicles (11). During pregnancy, the placenta releases vesicles such as syncytial nuclear aggregates, macrovesicles, nanovesicles, and exosomes (48). Studies have shown that pregnancies with obstetric complications have differences in cargo and size of vesicles released from trophoblast cells (34). To investigate if mtDNA from trophoblast cells is sequestered into extracellular vesicles when released from trophoblast cells, we quantified mtDNA copy number in EV-BeWo and found an increase in mtDNA levels stored in EV-BeWo exposed to ROS. This finding indicates that trophoblast cells release EVs enriched in mtDNA in response to ROS and cell death. However, we do not exclude the possibility that mtDNA is also contained in other vesicular structures derived from these cells such as liposomes or apoptotic bodies. The characterization of EVs in our study (i.e., NTA) indicate, however, there are not any apoptotic bodies present in our samples as there was no additional peak and no signal showing at sizes >500 nm (49). Our data establishes the foundation for future investigations to use markers of EV-containing mtDNA to establish cellular origin of ccf-mtDNA in an *in vivo* model.

Previous studies have implicated the role of autophagy impairment in placental pathology associated with ROS (50–56). Studies in trophoblast cells demonstrate an activation of autophagy after exposure to hypoxia(54, 55) or pharmacological inducers of ROS(_38_) with increased expression of LC3 and decreased p62 expression. LC3 is a marker of autophagosome formation and p62 is a cargo loading protein (57). Similarly, in this study we demonstrate that ROS potentially increased autophagy as suggested by an increase in LC3 and a reduction in p62. We hypothesize that ROS mediates the balance between autophagy and cell death pathways with autophagy functioning as a cytoprotective mechanism until high levels of ROS initiate cell death and release of mtDNA (52, 54, 58). Additional studies are needed to investigate the role of autophagy in mtDNA release and its role in placental pathology. Moreover, other mechanisms, including mitophagy (54, 59, 60) and anti-oxidant production (35, 58), may also be implicated as cytoprotective mechanisms in trophoblast cell responses to ROS and their relationship to mtDNA release warrants further investigation (35, 61).

### Conclusions, Limitations, and Future Directions

The main objective of the present study was to establish an association between oxidative stress, which is a common feature of placental pathology in obstetric complications, and mtDNA release mechanisms. We tested our hypothesis using a commonly used *in vitro* model of trophoblast cells, the BeWo choriocarcinoma monolayer cell line. Proteomics analysis has shown that the protein profile of BeWo cells resembles that of villous trophoblasts(62). This cell line has been widely used since the 1980s as a surrogate of villous trophoblast cells(63). Although primary trophoblast cells would an ideal choice, they can be used only for short term experiments due to their inability to proliferate in culture(64). A limitation of using BeWo cells is that we cannot address the effects of gestational age on ROS-mediated release of mtDNA or the interaction of trophoblast cells with other cell types (i.e., endothelial cells) and thus, future experiments in placental explains or organoids may be necessary to validate our findings. It is noteworthy that BeWo cells can spontaneously syncytialize although their basal fusion rate, in the absence of an inducer, is relatively low as compared to other choriocarcinoma cell lines (64). Nevertheless, due to their ability to spontaneously fuse to form syncytia, BeWo cells may express both cytotrophoblast and syncytiotrophoblast markers. In our flow cytometry experiments, we noted the presence of two distinct cell populations of BeWo cells. This heterogeneity may reflect syncytialized and non-syncytialized cells; however, we did not specifically characterize these. Similar data have been previously reported by other investigators (20). Future studies should examine the propensity of various trophoblast cell types to release mtDNA in response to stress using primary trophoblast cells or other *in vitro* models such as placental explants. In the present investigation, we measured mtDNA content in BeWo-derived EVs based on our findings that most ccf-mtDNA(10) as well as extracellular mtDNA in cell cultures (current study) is membrane-bound. In the EV characterization data, we noted expression of GM130, a marker of the Golgi apparatus. This indicates that there could be apoptotic bodies present in samples; however, as mentioned previously, the NTA data do not show a signal at >500+ nm size (65). This finding might also reflect contamination from non-exosomal proteins due to co-precipitation as a result of the polymer-based precipitation technique utilized to isolate exosomes (66).

In conclusion, our results demonstrated that BeWo cells, a well-established model of trophoblast cells, release mtDNA constitutively and this release is enhanced in response to ROS. Further, we established an association between ROS, mtDNA release, autophagy, and non-apoptotic cell death mechanisms. The current investigation reveals molecular targets and potential mechanisms that can be used in future investigations to validate our main outcomes in non-immortalized trophoblast cells, and to investigate the source and biomarker potential of ccf-mtDNA in *in vivo* preclinical models and humans.

## ACKNOWLEDGEMENTS

We acknowledge Sharad Shrestha, Ph.D. in the University of North Texas Health Science Center at Fort Worth Research Core Labs for training and assistance with the generation of Flow Cytometry data.

Present address of Spencer. C. Cushen: Department of Surgery, Virginia Mason Medical Center, Seattle, WA, USA.

## Grants

This study was supported by NIH R01 HL146562 and HL146562-S2 to SG, AHA 18PRE33960162 and NIH T32 AG020494 to SCC, AHA 19TPA-34850131 to SG, AHA 22POST-903250 to JB, AHA 22PRE-900431 to JG, AHA 23PRE-1012811 to ST. The content is solely the responsibility of the authors and does not necessarily represent the official views of the National Institutes of Health.

## DISCLOSURES

No conflicts of interest, financial or otherwise, are declared by the author(s).

## AUTHOR CONTRIBUTIONS

JG, SCC, and SG designed the study and experiments. JG, SCC, IG, RNO, ST, and JB collected data. JG, SCC, NH, ZZ, IG, SG, and NRP analyzed data. JG, NRP, ZZ, and SG performed statistical analysis. All authors contributed to data interpretation. JG, SCC, RNO, NH, and SG prepared figures. JG and SG drafted manuscript. All authors edited manuscript. All authors approved the final version of the manuscript.

## Notes

### Competing Interest Statement

The authors have declared no competing interest.

## REFERENCES

1. Soares MJ, Varberg KM, and Iqbal K. Hemochorial placentation: development, function, and adaptations. Biol Reprod 99: 196–211, 2018.

2. Redman CWG, Staff AC, and Roberts JM. Syncytiotrophoblast stress in preeclampsia: the convergence point for multiple pathways. Am J Obstet Gynecol 226: S907–S927, 2022.

3. Diaz P, Powell TL, and Jansson T. The role of placental nutrient sensing in maternal-fetal resource allocation. Biol Reprod 91: 82, 2014.

4. Gude NM, Roberts CT, Kalionis B, and King RG. Growth and function of the normal human placenta. Thromb Res 114: 397–407, 2004.

5. West AP, and Shadel GS. Mitochondrial DNA in innate immune responses and inflammatory pathology. Nat Rev Immunol 17: 363–375, 2017.

6. Cushen SC, Sprouse ML, Blessing A, Sun J, Jarvis SS, Okada Y, Fu Q, Romero SA, Phillips NR, and Goulopoulou S. Cell-free mitochondrial DNA increases in maternal circulation during healthy pregnancy: a prospective, longitudinal study. Am J Physiol Regul Integr Comp Physiol 318: R445–R452, 2020.

7. Qiu C, Hevner K, Enquobahrie DA, and Williams MA. A case-control study of maternal blood mitochondrial DNA copy number and preeclampsia risk. Int J Mol Epidemiol Genet 3: 237–244, 2012.

8. Marschalek J, Wohlrab P, Ott J, Wojta J, Speidl W, Klein KU, Kiss H, Pateisky P, Zeisler H, and Kuessel L. Maternal serum mitochondrial DNA (mtDNA) levels are elevated in preeclampsia - A matched case-control study. Pregnancy Hypertens 14: 195–199, 2018.

9. Busnelli A, Lattuada D, Ferrari S, Reschini M, Colciaghi B, Somigliana E, Fedele L, and Ferrazzi E. Mitochondrial DNA Copy Number in Peripheral Blood in the First Trimester of Pregnancy and Different Preeclampsia Clinical Phenotypes Development: A Pilot Study. Reprod Sci 26: 1054–1061, 2019.

10. Cushen SC, Ricci CA, Bradshaw JL, Silzer T, Blessing A, Sun J, Zhou Z, Scroggins SM, Santillan MK, Santillan DA, Phillips NR, and Goulopoulou S. Reduced Maternal Circulating Cell-Free Mitochondrial DNA Is Associated With the Development of Preeclampsia. J Am Heart Assoc 11: e021726, 2022.

11. Trumpff C, Michelson J, Lagranha CJ, Taleon V, Karan KR, Sturm G, Lindqvist D, Fernstrom J, Moser D, Kaufman BA, and Picard M. Stress and circulating cell-free mitochondrial DNA: A systematic review of human studies, physiological considerations, and technical recommendations. Mitochondrion 59: 225–245, 2021.

12. Wiersma M, van Marion DMS, Bouman EJ, Li J, Zhang D, Ramos KS, Lanters EAH, de Groot NMS, and Brundel B. Cell-Free Circulating Mitochondrial DNA: A Potential Blood-Based Marker for Atrial Fibrillation. Cells 9: 2020.

13. Nakahira K, Kyung SY, Rogers AJ, Gazourian L, Youn S, Massaro AF, Quintana C, Osorio JC, Wang Z, Zhao Y, Lawler LA, Christie JD, Meyer NJ, Mc Causland FR, Waikar SS, Waxman AB, Chung RT, Bueno R, Rosas IO, Fredenburgh LE, Baron RM, Christiani DC, Hunninghake GM, and Choi AM. Circulating mitochondrial DNA in patients in the ICU as a marker of mortality: derivation and validation. PLoS Med 10: e1001577; discussion e1001577, 2013.

14. Meng X, Schwarzenbach H, Yang Y, Muller V, Li N, Tian D, Shen Y, and Gong Z. Circulating Mitochondrial DNA is Linked to Progression and Prognosis of Epithelial Ovarian Cancer. Transl Oncol 12: 1213–1220, 2019.

15. De Gaetano A, Solodka K, Zanini G, Selleri V, Mattioli AV, Nasi M, and Pinti M. Molecular Mechanisms of mtDNA-Mediated Inflammation. Cells 10: 2021.

16. Nakahira K, Haspel JA, Rathinam VA, Lee SJ, Dolinay T, Lam HC, Englert JA, Rabinovitch M, Cernadas M, Kim HP, Fitzgerald KA, Ryter SW, and Choi AM. Autophagy proteins regulate innate immune responses by inhibiting the release of mitochondrial DNA mediated by the NALP3 inflammasome. Nat Immunol 12: 222–230, 2011.

17. Kim J, Gupta R, Blanco LP, Yang S, Shteinfer-Kuzmine A, Wang K, Zhu J, Yoon HE, Wang X, Kerkhofs M, Kang H, Brown AL, Park SJ, Xu X, Zandee van Rilland E, Kim MK, Cohen JI, Kaplan MJ, Shoshan-Barmatz V, and Chung JH. VDAC oligomers form mitochondrial pores to release mtDNA fragments and promote lupus-like disease. Science 366: 1531–1536, 2019.

18. Ives CW, Sinkey R, Rajapreyar I, Tita ATN, and Oparil S. Preeclampsia-Pathophysiology and Clinical Presentations: JACC State-of-the-Art Review. J Am Coll Cardiol 76: 1690–1702, 2020.

19. King A, Thomas L, and Bischof P. Cell culture models of trophoblast II: trophoblast cell lines--a workshop report. Placenta 21 Suppl A: S113–119, 2000.

20. Khera A, Vanderlelie JJ, Holland O, and Perkins AV. Overexpression of Endogenous Anti-Oxidants with Selenium Supplementation Protects Trophoblast Cells from Reactive Oxygen Species-Induced Apoptosis in a Bcl-2-Dependent Manner. Biol Trace Elem Res 177: 394–403, 2017.

21. Osikoya O, Cushen SC, Gardner JJ, Raetz MM, Nagarajan B, Raut S, and Goulopoulou S. Exosomes facilitate intercellular communication between uterine perivascular adipose tissue and vascular smooth muscle cells in pregnant rats. Am J Physiol Heart Circ Physiol 323: H577–H584, 2022.

22. Phillips NR, Sprouse ML, and Roby RK. Simultaneous quantification of mitochondrial DNA copy number and deletion ratio: a multiplex real-time PCR assay. Sci Rep 4: 3887, 2014.

23. Lima K, de Miranda LBL, Del Milagro Bernabe Garnique A, de Almeida BO, do Nascimento MC, Alcantara GAS, Machado-Santelli GM, Rego EM, and Machado-Neto JA. The Multikinase Inhibitor AD80 Induces Mitotic Catastrophe and Autophagy in Pancreatic Cancer Cells. Cancers (Basel) 15: 2023.

24. Cohen J. *Statistical power analysis for the behavioral sciences*. Hillsdale, N.J.: L. Erlbaum Associates, 1988, p. xxi, 567 pages.

25. Boudreau LH, Duchez AC, Cloutier N, Soulet D, Martin N, Bollinger J, Pare A, Rousseau M, Naika GS, Levesque T, Laflamme C, Marcoux G, Lambeau G, Farndale RW, Pouliot M, Hamzeh-Cognasse H, Cognasse F, Garraud O, Nigrovic PA, Guderley H, Lacroix S, Thibault L, Semple JW, Gelb MH, and Boilard E. Platelets release mitochondria serving as substrate for bactericidal group IIA-secreted phospholipase A2 to promote inflammation. Blood 124: 2173–2183, 2014.

26. Thurairajah K, Briggs GD, and Balogh ZJ. The source of cell-free mitochondrial DNA in trauma and potential therapeutic strategies. Eur J Trauma Emerg Surg 44: 325–334, 2018.

27. Hayakawa K, Chan SJ, Mandeville ET, Park JH, Bruzzese M, Montaner J, Arai K, Rosell A, and Lo EH. Protective Effects of Endothelial Progenitor Cell-Derived Extracellular Mitochondria in Brain Endothelium. Stem Cells 36: 1404–1410, 2018.

28. Stigliani S, Orlando G, Massarotti C, Casciano I, Bovis F, Anserini P, Ubaldi FM, Remorgida V, Rienzi L, and Scaruffi P. Non-invasive mitochondrial DNA quantification on Day 3 predicts blastocyst development: a prospective, blinded, multi-centric study. Mol Hum Reprod 25: 527–537, 2019.

29. Munakata Y, Shirasuna K, Kuwayama T, and Iwata H. Cell-free DNA in medium is associated with the maturation ability of in vitro cultured oocytes. J Reprod Dev 65: 171–175, 2019.

30. Al Amir Dache Z, Otandault A, Tanos R, Pastor B, Meddeb R, Sanchez C, Arena G, Lasorsa L, Bennett A, Grange T, El Messaoudi S, Mazard T, Prevostel C, and Thierry AR. Blood contains circulating cell-free respiratory competent mitochondria. FASEB J 34: 3616–3630, 2020.

31. Song X, Hu W, Yu H, Wang H, Zhao Y, Korngold R, and Zhao Y. Existence of Circulating Mitochondria in Human and Animal Peripheral Blood. Int J Mol Sci 21: 2020.

32. Lin HY, Liou CW, Chen SD, Hsu TY, Chuang JH, Wang PW, Huang ST, Tiao MM, Chen JB, Lin TK, and Chuang YC. Mitochondrial transfer from Wharton’s jelly-derived mesenchymal stem cells to mitochondria-defective cells recaptures impaired mitochondrial function. Mitochondrion 22: 31–44, 2015.

33. Colleoni F, Lattuada D, Garretto A, Massari M, Mando C, Somigliana E, and Cetin I. Maternal blood mitochondrial DNA content during normal and intrauterine growth restricted (IUGR) pregnancy. Am J Obstet Gynecol 203: 365 e361–366, 2010.

34. Tong M, Johansson C, Xiao F, Stone PR, James JL, Chen Q, Cree LM, and Chamley LW. Antiphospholipid antibodies increase the levels of mitochondrial DNA in placental extracellular vesicles: Alarmin-g for preeclampsia. Sci Rep 7: 16556, 2017.

35. Vaka VR, McMaster KM, Cunningham MW, Jr., Ibrahim T, Hazlewood R, Usry N, Cornelius DC, Amaral LM, and LaMarca B. Role of Mitochondrial Dysfunction and Reactive Oxygen Species in Mediating Hypertension in the Reduced Uterine Perfusion Pressure Rat Model of Preeclampsia. Hypertension 72: 703–711, 2018.

36. Sedeek M, Gilbert JS, LaMarca BB, Sholook M, Chandler DL, Wang Y, and Granger JP. Role of reactive oxygen species in hypertension produced by reduced uterine perfusion in pregnant rats. Am J Hypertens 21: 1152–1156, 2008.

37. Zhao RZ, Jiang S, Zhang L, and Yu ZB. Mitochondrial electron transport chain, ROS generation and uncoupling (Review). Int J Mol Med 44: 3–15, 2019.

38. Heazell AE, Taylor NN, Greenwood SL, Baker PN, and Crocker IP. Does altered oxygenation or reactive oxygen species alter cell turnover of BeWo choriocarcinoma cells? Reprod Biomed Online 18: 111–119, 2009.

39. Vangrieken P, Remels AHV, Al-Nasiry S, Bast A, Janssen GMJ, von Rango U, Vroomans D, Pinckers YCW, van Schooten FJ, and Schiffers PMH. Placental hypoxia-induced alterations in vascular function, morphology, and endothelial barrier integrity. Hypertens Res 43: 1361–1374, 2020.

40. van der Pol A, van Gilst WH, Voors AA, and van der Meer P. Treating oxidative stress in heart failure: past, present and future. Eur J Heart Fail 21: 425–435, 2019.

41. Wang Y, Zhang SX, and Gozal D. Reactive oxygen species and the brain in sleep apnea. Respir Physiol Neurobiol 174: 307–316, 2010.

42. Walker OS, Ragos R, Wong MK, Adam M, Cheung A, and Raha S. Reactive oxygen species from mitochondria impacts trophoblast fusion and the production of endocrine hormones by syncytiotrophoblasts. PLoS One 15: e0229332, 2020.

43. Al-Nasiry S, Spitz B, Hanssens M, Luyten C, and Pijnenborg R. Differential effects of inducers of syncytialization and apoptosis on BeWo and JEG-3 choriocarcinoma cells. Hum Reprod 21: 193–201, 2006.

44. Forkink M, Basit F, Teixeira J, Swarts HG, Koopman WJH, and Willems P. Complex I and complex III inhibition specifically increase cytosolic hydrogen peroxide levels without inducing oxidative stress in HEK293 cells. Redox Biol 6: 607–616, 2015.

45. Siddiqui MA, Ahmad J, Farshori NN, Saquib Q, Jahan S, Kashyap MP, Ahamed M, Musarrat J, and Al-Khedhairy AA. Rotenone-induced oxidative stress and apoptosis in human liver HepG2 cells. Mol Cell Biochem 384: 59–69, 2013.

46. Zuberek M, Paciorek P, Rakowski M, and Grzelak A. How to Use Respiratory Chain Inhibitors in Toxicology Studies-Whole-Cell Measurements. Int J Mol Sci 23: 2022.

47. Newman LE, and Shadel GS. Mitochondrial DNA Release in Innate Immune Signaling. Annu Rev Biochem 92: 299–332, 2023.

48. Tong M, and Chamley LW. Placental extracellular vesicles and feto-maternal communication. Cold Spring Harb Perspect Med 5: a023028, 2015.

49. Paul N, Sultana Z, Fisher JJ, Maiti K, and Smith R. Extracellular vesicles-crucial players in human pregnancy. Placenta 140: 30–38, 2023.

50. Hung TH, Chen SF, Lo LM, Li MJ, Yeh YL, and Hsieh TT. Increased autophagy in placentas of intrauterine growth-restricted pregnancies. PLoS One 7: e40957, 2012.

51. Oh SY, Choi SJ, Kim KH, Cho EY, Kim JH, and Roh CR. Autophagy-related proteins, LC3 and Beclin-1, in placentas from pregnancies complicated by preeclampsia. Reprod Sci 15: 912–920, 2008.

52. Gao L, Qi HB, Kamana KC, Zhang XM, Zhang H, and Baker PN. Excessive autophagy induces the failure of trophoblast invasion and vasculature: possible relevance to the pathogenesis of preeclampsia. J Hypertens 33: 106–117, 2015.

53. Akaishi R, Yamada T, Nakabayashi K, Nishihara H, Furuta I, Kojima T, Morikawa M, Yamada T, Fujita N, and Minakami H. Autophagy in the placenta of women with hypertensive disorders in pregnancy. Placenta 35: 974–980, 2014.

54. Vangrieken P, Al-Nasiry S, Bast A, Leermakers PA, Tulen CBM, Janssen GMJ, Kaminski I, Geomini I, Lemmens T, Schiffers PMH, van Schooten FJ, and Remels AHV. Hypoxia-induced mitochondrial abnormalities in cells of the placenta. PLoS One 16: e0245155, 2021.

55. Chen B, Longtine MS, and Nelson DM. Hypoxia induces autophagy in primary human trophoblasts. Endocrinology 153: 4946–4954, 2012.

56. Cheng S, Huang Z, Jash S, Wu K, Saito S, Nakashima A, and Sharma S. Hypoxia-Reoxygenation Impairs Autophagy-Lysosomal Machinery in Primary Human Trophoblasts Mimicking Placental Pathology of Early-Onset Preeclampsia. Int J Mol Sci 23: 2022.

57. Yoshii SR, and Mizushima N. Monitoring and Measuring Autophagy. Int J Mol Sci 18: 2017.

58. Wang P, Huang CX, Gao JJ, Shi Y, Li H, Yan H, Yan SJ, and Zhang Z. Resveratrol induces SIRT1-Dependent autophagy to prevent H(2)O(2)-Induced oxidative stress and apoptosis in HTR8/SVneo cells. Placenta 91: 11–18, 2020.

59. Sun Y, Lv D, Xie Y, Xu H, Li X, Li F, Fan Y, Zhang X, Zhang Y, Chen S, He M, and Deng D. PINK1-mediated mitophagy induction protects against preeclampsia by decreasing ROS and trophoblast pyroptosis. Placenta 143: 1–11, 2023.

60. Zhou X, Zhao X, Zhou W, Qi H, Zhang H, Han TL, and Baker P. Impaired placental mitophagy and oxidative stress are associated with dysregulated BNIP3 in preeclampsia. Sci Rep 11: 20469, 2021.

61. Ausman J, Abbade J, Ermini L, Farrell A, Tagliaferro A, Post M, and Caniggia I. Ceramide-induced BOK promotes mitochondrial fission in preeclampsia. Cell Death Dis 9: 298, 2018.

62. Szklanna PB, Wynne K, Nolan M, Egan K, Ainle FN, and Maguire PB. Comparative proteomic analysis of trophoblast cell models reveals their differential phenotypes, potential uses, and limitations. Proteomics 17: e1700037, 2017.

63. Li X, Li ZH, Wang YX, and Liu TH. A comprehensive review of human trophoblast fusion models: recent developments and challenges. Cell Death Discov 9: 372, 2023.

64. Orendi K, Kivity V, Sammar M, Grimpel Y, Gonen R, Meiri H, Lubzens E, and Huppertz B. Placental and trophoblastic in vitro models to study preventive and therapeutic agents for preeclampsia. Placenta 32 Suppl: S49–54, 2011.

65. Keerthikumar S, Gangoda L, Liem M, Fonseka P, Atukorala I, Ozcitti C, Mechler A, Adda CG, Ang CS, and Mathivanan S. Proteogenomic analysis reveals exosomes are more oncogenic than ectosomes. Oncotarget 6: 15375–15396, 2015.

66. Li M, Lou D, Chen J, Shi K, Wang Y, Zhu Q, Liu F, and Zhang Y. Deep dive on the proteome of salivary extracellular vesicles: comparison between ultracentrifugation and polymer-based precipitation isolation. Anal Bioanal Chem 413: 365–375, 2021.

