## Supplemental Materials for "Oxidative stress induces release of mitochondrial DNA into the extracellular space in human placental villous trophoblast BeWo cells"

[1]Department of Physiology and Anatomy, University of North Texas Health Science Center, Fort Worth, Texas, USA; [2]Texas College of Osteopathic Medicine, University of North Texas Health Science Center, Fort Worth, Texas, USA; [3]Department of Microbiology, Immunology, and Genetics, University of North Texas Health Science Center, Fort Worth, Texas, USA; [4]Department of Population & Community Health, University of North Texas Health Science Center, Fort Worth, Texas, USA; [5]Department of Pharmaceutical Sciences in System College of Pharmacy, University of North Texas Health Science Center, Fort Worth, Texas, USA; [6]Lawrence D. Longo, MD Center for Perinatal Biology, Departments of Basic Sciences, Gynecology and Obstetrics, Loma Linda University School of Medicine, Loma Linda, CA, USA

**Methods**

Immunocytochemistry

BeWo cells were seeded at 80,000 cells/well on coverslips in a 24 well plate. Cells were grown to approximately 60% confluency, fixed with 4% paraformaldehyde (PFA) for 30 minutes, and incubated with blocking buffer (PBS, 10% goat serum, 0.25% Triton-X 100) for 1.5 hr. Slides were incubated with rabbit monoclonal anti-cytokeratin VII antibody (abcam, cat# ab181598, company location) or mouse monoclonal alpha smooth muscle antibody (Sigma, cat# A2547) for 30 min at room temperature. This was followed by incubation with fluorescence secondary antibodies, goat anti-rabbit IgG (abcam, ab150077) or goat anti-mouse IgG (abcam, cat# ab150113) for 1.5 hours. Coverslips were mounted to microscope slides using mounting media containing 4′,6-diamidino-2-phenylindole (DAPI; ProLong Diamond, Thermo, cat# P36966) to visualize nuclei. Images were captured with an Olympus BX41 fluorescent microscope at 4x magnification and were exposed for 2 seconds (Alexa fluor 488) or auto exposed (DAPI) at a gain of ISO 400. Images captured at 40x magnification were captured using auto exposure at a gain of ISO 400.

### Figures

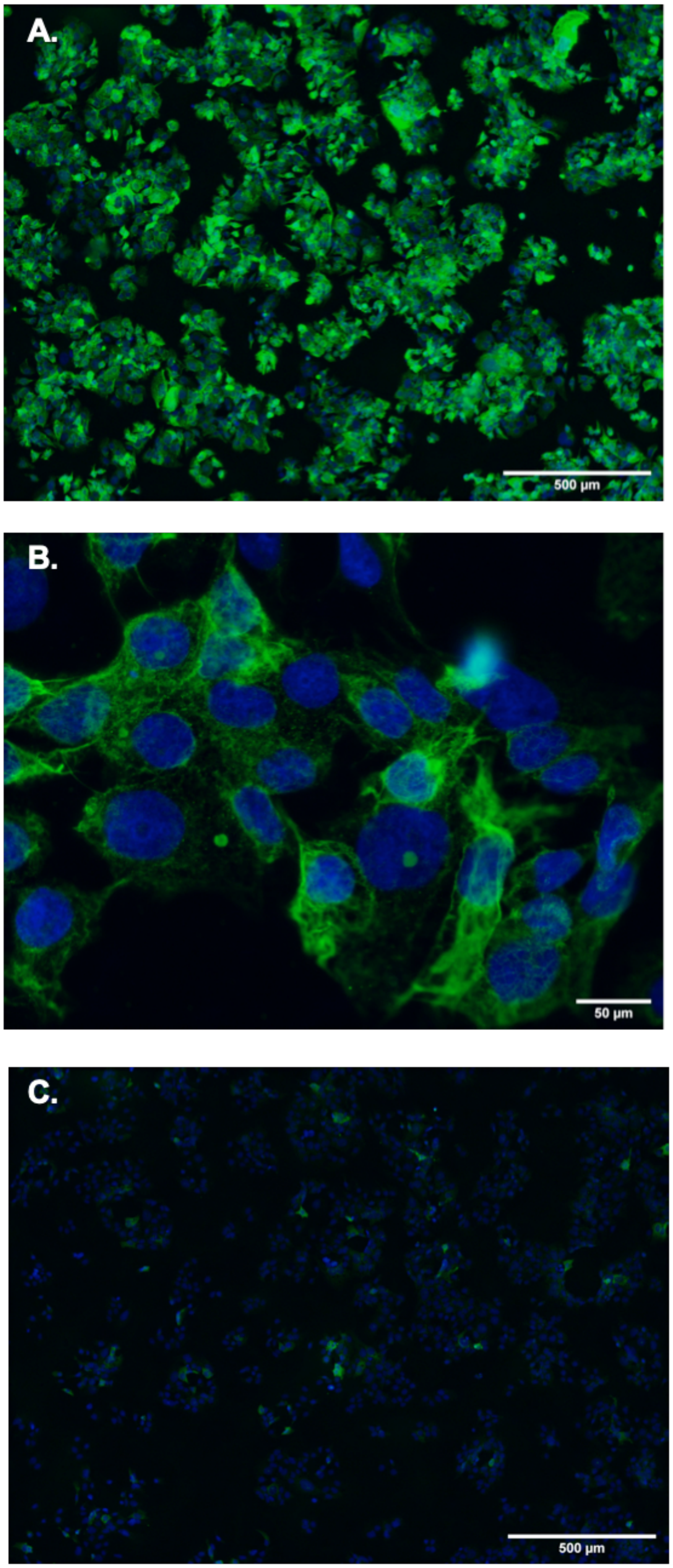

**Supplementary Figure 1— BeWo trophoblast cells characterization.** To characterize BeWo cells they were stained for cytokeratin VII (a marker of epithelial cells; green) and DAPI (a nuclear marker; blue) at A) 4x and B) 40x magnification. BeWo cells stained with α-smooth muscle actin (negative control; green) and the nuclear stain DAPI (blue) at C) 4x magnification.

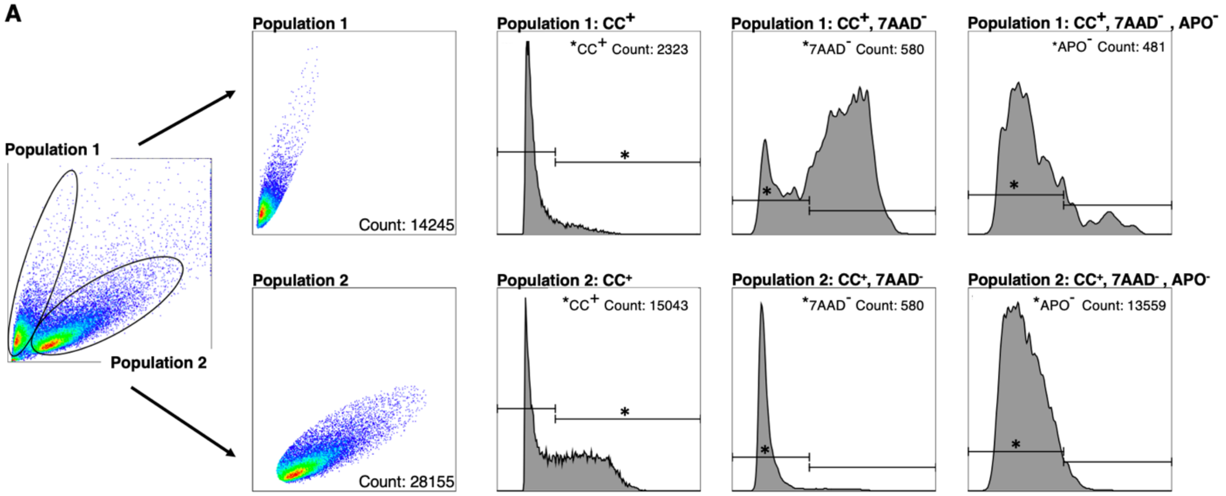


**B** $$\textbf{\% Viability} = \frac{\text{[Pop 1 (CC}^{+}\text{, 7AAD}^{-}\text{, APO}^{-}\text{) + Pop 2 (CC}^{+}\text{, 7AAD}^{-}\text{, APO}^{-}\text{) Counts]}}{\text{[Total Cell Count (Pop 1 + Pop 2)]}} \times 100$$

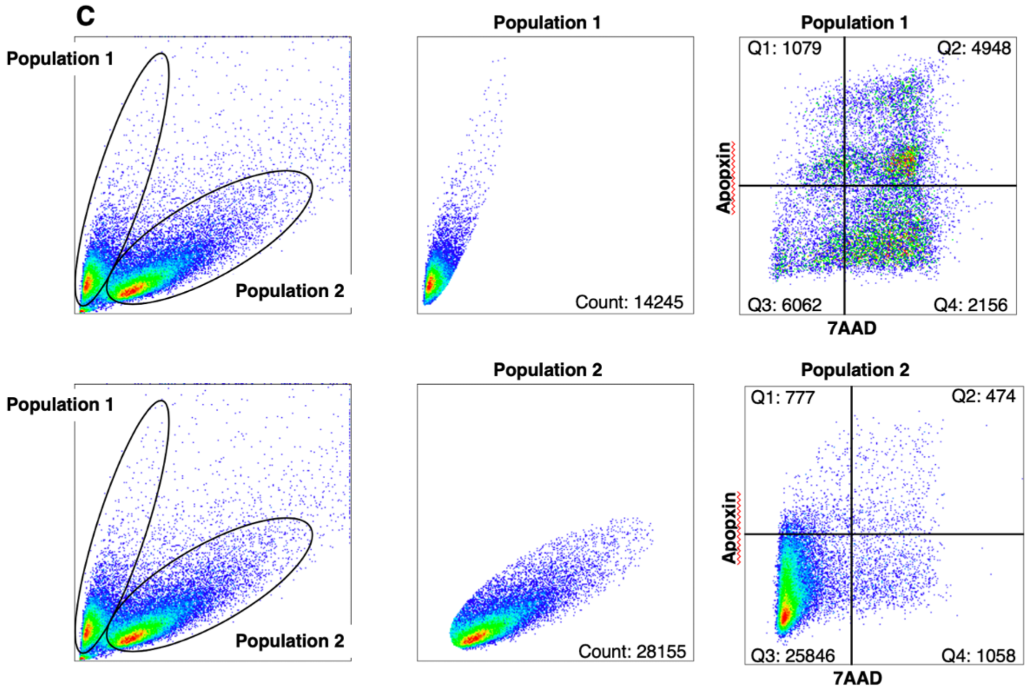

**D** **% Apoptosis=** $\frac{\text{[Pop 1 (Q1 + Q2) + Pop 2 (Q1 + Q2) Counts]}}{\text{[Total Cell Count (Pop 1 + Pop 2)]}} \times 100$

**E** **% Necrosis=** $\frac{\text{[Pop 1 (Q4) + Pop 2 (Q4) Counts]}}{\text{[Total Cell Count (Pop1 + Pop2)]}} \times 100$

**Supplementary Figure 2— Gating strategy for cell viability and cell death in BeWo trophoblast cells in response to oxidative stress.** Two populations of cells were identified, and gating strategies were applied to each population. Histograms were used to determine A) cell viability by gating CC+, 7AAD-, and Apo- cells and B) the percentage of viable cells was determined by taking CC+, 77AD-, Apo- cell counts and dividing by total number of cell counts. C) Apoptosis (Q1+Q2) and necrosis (Q3) were determined with 7AAD on the x-axis and Apopoxin on the y-axis. D) The percentage of total apoptosis was calculated by summing Q1 and Q3 counts and dividing by the total number of cell counts. E) The percentage of total necrosis was determined by dividing Q4 by the total number of cell counts.

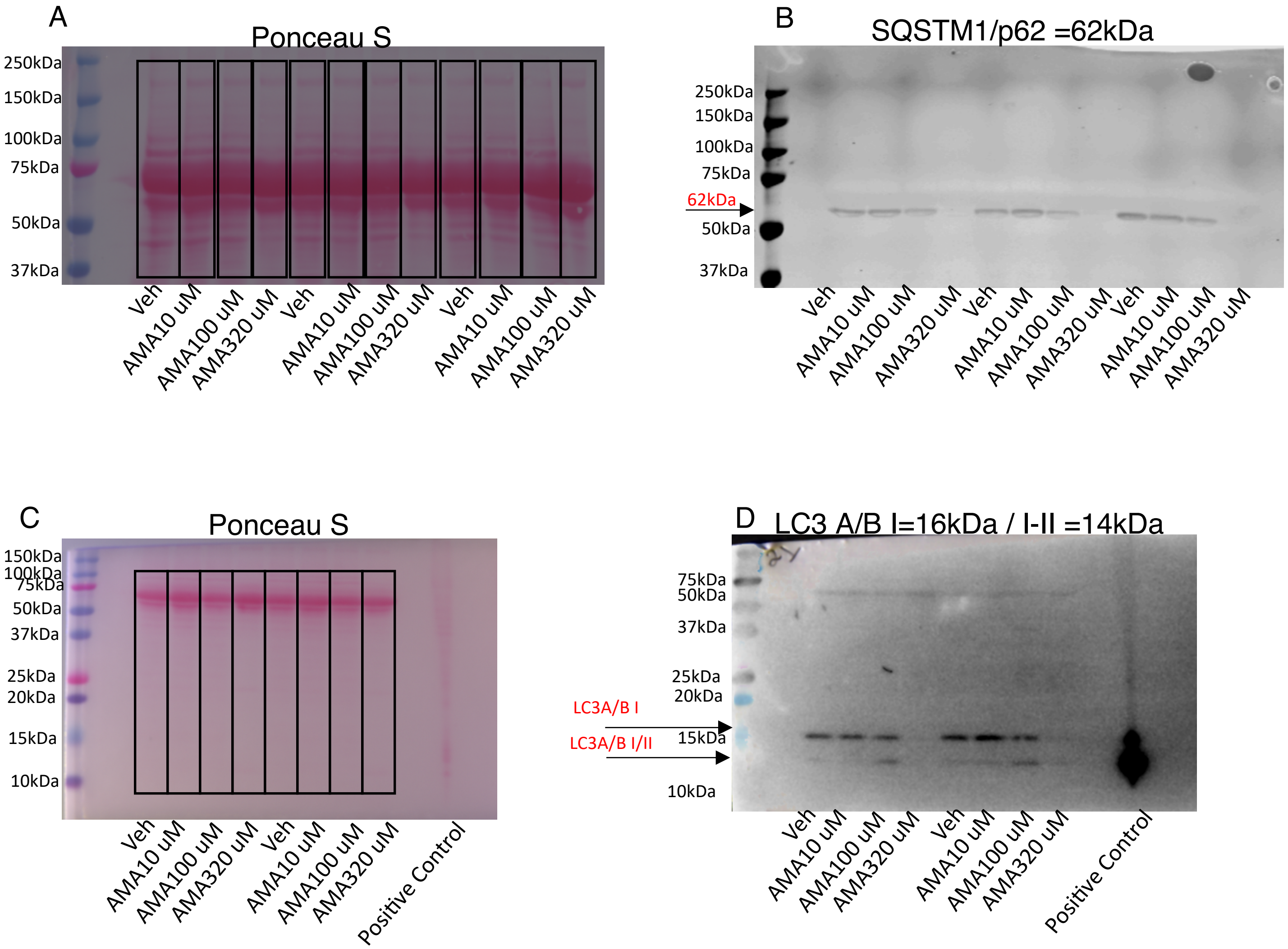


**Supplementary Figure 3**— Representative immunoblot membranes of SQSTM1/p62 (A-B) and LC3 (C-D) with respective Ponceau Stain. Area of analysis of total protein is indicated with rectangular boxes (A, C).

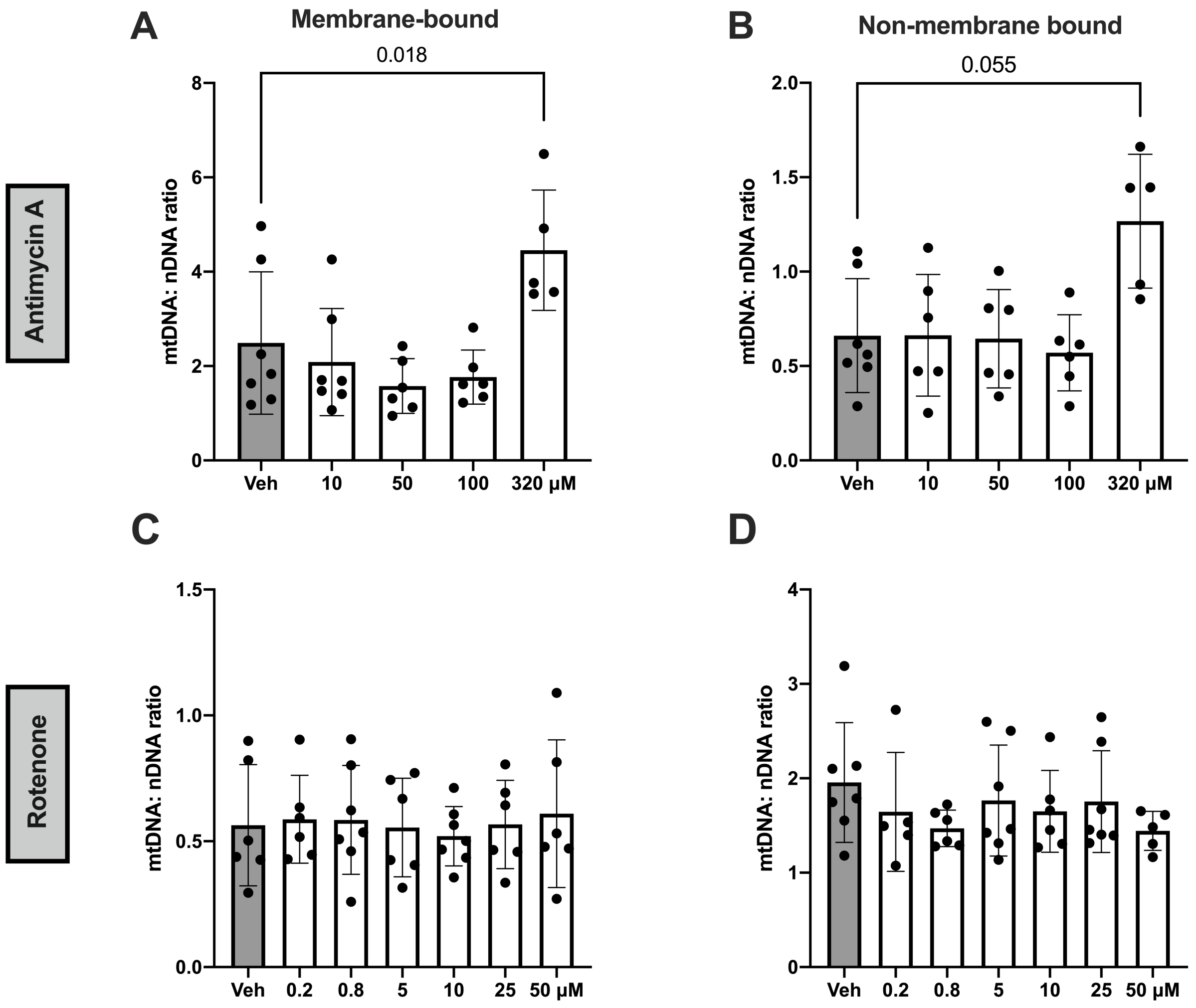


**Supplementary Figure 4— Effect of antimycin A and rotenone on mtDNA to nuclear DNA ratios in BeWo trophoblast cells.** The ratio of mtDNA to nDNA was increased in BeWo cells treated with antimycin A in both the A) membrane-bound (p=0.0016) and B) non-membrane bound forms (p=0.0041). Rotenone did not affect these ratios in the C) membrane-bound (p=0.64) or D) non-membrane bound form (p=0.99). These data were analyzed with one-way ANOVA (A, C) or Kruskal Wallis test (B, D) for data that were not normally distributed. Means ± SD. n=5-7 independent observations.

**Supplementary Table 1.** Chemical and reagents

| Chemical | Company | Catalog Number | City |
|---|---|---|---|
| Dimethylsulfoxide (DMSO) | ATCC | 4-X | Manassas, VA, USA |
| Tris | BioRad | 161-0716 | Hercules, CA, USA |
| Phenol Free Media | Gibco | 11039-021 | Grand Island, NY, USA |
| 2',7'-dichlorofluorescin diacetate (DCF-DA) | Invitrogen | D399 | Eugene, OR, USA |
| Antimycin | Sigma Aldrich | A8674 | Sigma, St. Louis, MO, USA |
| N-propyl gallate | Sigma Aldrich | 02370 | Sigma, St. Louis, MO, USA |
| Rotenone | Sigma Aldrich | R8875 | Sigma, St. Louis, MO, USA |
| Tert-butyl hydrogen peroxide | Sigma Aldrich | 458139 | Sigma, St. Louis, MO, USA |
| Triton X-100 | Sigma Aldrich | X100 | Sigma, St. Louis, MO, USA |
| Ethanol Absolute, 200 Proof, for Molecular Biology | ThermoFisher | BP2818 | Fair Lawn, NJ, USA |
| Glycerol | Fisher Scientific | BP229-1 | Fair Lawn, NJ, USA |
| Phosphate Buffered Saline | Fisher | BP399 | Fair Lawn, NJ, USA |
